# Effective soil extraction method for cultivating previously uncultured soil bacteria

**DOI:** 10.1101/322735

**Authors:** Tuan Manh Nguyen, Chan Seo, Moongi Ji, Man-Jeong Paik, Seung-Woon Myung, Jaisoo Kim

**Affiliations:** Department of Life Science, College of Natural Sciences and Engineering, Kyonggi University, Suwon, Gyeonggi-Do 16227, Republic of Korea; Department of Chemistry, College of Natural Sciences and Engineering, Kyonggi University, Suwon, Gyeonggi-Do 16227, Republic of Korea; College of Pharmacy and Research Institute of Life and Pharmaceutical Sciences, Sunchon National University, Suncheon, Jeollanam-do 57922, Republic of Korea; Thai Nguyen University of Agriculture and Forestry, Quyet Thang, Thai Nguyen, Vietnam

**Keywords:** uncultured bacteria, cultivation, new soil extract (NSE), intensive soil extract medium (ISEM), new taxonomic candidates, low-molecular-weight organic substances (LMWOS), isolation, subculture

## Abstract

Here, a new medium named as intensive soil extract medium (ISEM) based on new soil extract (NSE) using 80% ethanol was used to efficiently isolate previously uncultured bacteria and new taxonomic candidates, which accounted for 49% and 55% of the total isolates examined (n=258), respectively. The new isolates were affiliating with seven phyla such as *Proteobacteria, Acidobacteria, Firmicutes, Actinobacteria, Verrucomicrobia, Planctomycetes*, and *Bacteroidetes*. The result of chemical analysis showed that NSE included more diverse components of low-molecular-weight organic substances than two conventional soil extracts using distilled water. Cultivation of previously uncultured bacteria is expected to extend knowledge through the discovery of new phenotypic, physiological and functional properties, and even roles of unknown genes.

**IMPORTANCE:** Either metagenomics or single-cell sequencing can detect unknown genes from uncultured microbial strains in environments and may find their significant potential metabolites and roles. However, such gene/genome-based techniques still have a critical problem making impossible for further applications through cultivation. To solve this problem, various approaches for cultivation of uncultured bacteria have been developed, but they still have lack of skill to grow them on solid media for isolation and subculture.

---

Approximately 1% of the total soil bacteria have been cultivated under laboratory conditions (1).Since the establishment of solid culture media, secondary metabolites have been isolated from microorganisms cultured in laboratories. Molecular tools revealed that prokaryotic species are very diverse and abundant in soil, and contain numerous unexplored potential metabolites (1–4). Although these tools enable analysis of the broad range of metabolic diversity of microorganisms without the need to isolate species, many bacterial characteristics are unknown because of limitations of cultivation (5, 6). The lack of complex factors/conditions in the laboratory has contributed to the inability to isolate various species (7, 8). Since the concept of “uncultured bacteria” was published in 1990 (9) to refer these bacteria as not yet cultured in laboratories, several methods have been developed in an attempt to culture these bacteria so far. These methods involved transporting bacteria from their natural environment to the laboratory for growth in artificial media/conditions similar to those in the natural environment by modifying growth media components (7) or growth conditions such pH and salt concentrations (9, 10), adding inorganic compounds or metals as electron donors/acceptors (11, 12), using various factors (13), coculture with helper bacteria (14, 15), soil extracts using water (16, 17) or aqueous buffers (18,>19), diluted medium or serial dilution culture (20, 21), long incubation time (10, 22), etc. Furthermore, sophisticated techniques were developed such as iChip for *in situ* cultivation (23), micro-bioreactor (24), optical tweezers (25), or micro-manipulator (26), which allowed analysis of individual cells in soil samples. However, new artificial media to maintain these cultures are needed. Although scientists can enrich slow-growing microorganisms using diffusion chambers (27, 28) or soil substrate membranes (29), most enriched bacteria do not grow on agar plates for isolation and further cultivation. Without successful cultivation, it is difficult to detect and identify novel organisms, obtain phenotypic and functional information, and determine the functions of unknown genes (30). The most important factor affecting the cultivation of uncultured bacteria and the most appropriate media conditions remain unclear.

Here, we developed a simple culture method based on new soil extract (NSE) using 80% ethanol without specific supplies and successfully cultured many previously uncultured bacterial strains. To evaluate our method, we checked the proportion of uncultured strains among isolates as well as that of new taxa, and analysed chemical components of NSE to compare with two traditional soil extracts (TSEs) commonly used.

## RESULTS

### Composition of soil extracts utilized as culture supplements

For nutrition and bacterial growth, heterotrophic soil bacteria depend on low-molecular-weight organic substances (LMWOS) and inorganic compounds (31). Thus, in this study, we compared the major LMWOS such as amino acids, fatty acids, and organic acids and inorganic ions in three different extraction methods (NSE: new extraction developed in this study; TSE1: traditional extraction without autoclaving; TSE2: traditional extraction with autoclaving) (Table 1). To overcome these problems, we used a mixture of methanol/water (4:1; 80%) to extract bacterial nutrients from soil rather than water or aqueous buffers and named this extract as new soil extract (NSE). Briefly, 500 g dry soil was prepared and shaken at 150 r.p.m with 1.3 L 80% methanol overnight at room temperature. The supernatant was transferred to a new flask and fresh 1.3 L 80% methanol was added to the remaining soil and mixed well for 1 h. The two supernatants were combined, filtered, and evaporated. The pure NSE was stored at 4°C until use. For amino acids, NSE showed a total yield of 18.47 mgL^−1^, which is much higher than for the other two methods which showed values of 4.83 and 5.87 mgL^−1^, respectively (P<0.006). Additionally, 21 amino acids were extracted, including four more amino acids (valine, pipecolic acid, serine, and threonine) and a very high concentration of tyrosine (9.29 mgL^−1^) compared to the other methods. This higher concentration and diversity of many amino acids in the NSE than in other two methods may cause better cultivability of uncultured soil bacteria. However, there was no difference between methods TSE1 and TSE2 (P_TSE1 vs TSE2_>0.05), indicating that autoclaving at 121°C had no effects. The results of fatty acid analysis could not be compared because method NSE extracted a much higher total concentration (13.11 mgL^−1^ vs. 0.79 mgL^−1^) and greater number (n=16 vs. n=8) of fatty acids than other two (Table 1). Greater amounts and larger numbers of fatty acids may improve the cultivability of uncultured soil bacteria. For organic acids, the results showed that method NSE had lower total yield of organic acids (25.13 mgL^−1^ vs. 62.69-89.00 mgL^−1^; *P* <0.03) but a greater total number (n=16 vs. n=11) including lactic acid, glycolic acid, 2-hydroxybuyric acid, fumaric acid, and α-ketoglutaric acid (Table 1). The total amount of organic acids obtained by methods TSE1 and TSE2 were significantly influenced by two major components, acetoacetic acid and oxaloacetic acid, but these did not significantly affect the total amount for method NSE.

The total yields of each approach regardless of inorganic composition were 748.03, 965.00, and 946.91 mgL^−1^, respectively. Although method NSE gave a lower concentration of total inorganic compounds than the other methods, the methanol-water mixture appeared to dissolve substances similarly to water and recovered all tested inorganic ions as water extractions (methods TSE1 & TSE2) (Table 1). Autoclaving did not significantly affect the dissolution of inorganic or organic compounds in water between methods TSE1 and TSE2 (P>0.7).

**Table 1.** **List of soil extract components derived from a new way and two general approaches.**

| Ingredients (mgL<sup>-1</sup>) | Soil extraction methods |  |  |  |  |  |  |
| --- | --- | --- | --- | --- | --- | --- | --- |
|  | NSE | TSE1 | TSE2 |  | NSE | TSE1 | TSE2 |
| **1. Amino acid** |  |  |  | *(Continued)* |  |  |  |
| Alanine | 0.466 | 0.099 | 0.178 | Erucic acid | nd | 0.071 | 0.064 |
| Glycine | 0.318 | 0.156 | 0.399 | Behenic acid | 1.799 | nd | nd |
| Valine | 0.086 | 0.000 | 0.000 | Nervonic acid | 0.074 | nd | nd |
| Leucine | 0.468 | 0.052 | 0.061 | Lignoceric acid | 2.603 | 0.093 | 0.079 |
| Isoleucine | 0.201 | 0.026 | 0.034 | Cerotic acid | 0.094 | 0.182 | 0.144 |
| Proline | 0.214 | 0.631 | 0.694 | **Total** | **13.114** | **0.789** | **0.793** |
| γ-Aminobutric acid | 0.756 | 0.621 | 0.689 | **3. Organic acid** |  |  |  |
| Pipecolic acid | 0.092 | nd | nd | 3-Hydroxybutyric acid | 0.962 | 2.068 | 1.881 |
| Pyroglutamic acid | 0.847 | 0.234 | 0.318 | Pyruvic acid | 0.404 | 4.310 | 5.746 |
| Serine | 0.379 | nd | nd | Acetoacetic acid | 1.465 | 28.750 | 27.247 |
| Threonine | 1.379 | nd | nd | Lactic acid | 9.418 | nd | nd |
| Phenylalanine | 0.219 | 0.033 | 0.036 | Glycolic acid | 8.031 | nd | nd |
| Cysteine | 0.216 | 0.117 | 0.293 | 2-Hydroxybutyric acid | 0.094 | nd | nd |
| Aspartic acid | 0.272 | 0.076 | 0.308 | Malonic acid | 0.845 | 1.205 | 1.754 |
| Glutamic acid | 0.552 | 0.398 | 0.392 | Succinic acid | 1.425 | 2.176 | 1.784 |
| Asparagine | 0.565 | 0.409 | 0.518 | Fumaric acid | 0.051 | nd | nd |
| Ornithine | 0.766 | 0.463 | 0.454 | Oxaloacetic acid | 0.173 | 18.353 | 44.839 |
| Glutamine | 0.547 | 0.668 | 0.664 | α-Ketoglutaric acid | 0.114 | nd | nd |
| Lysine | 0.122 | 0.246 | 0.235 | Malic acid | 0.728 | 3.322 | 3.332 |
| Tyrosine | 9.286 | 0.100 | 0.085 | 2-Hydroxyglutaric acid | 0.496 | 0.932 | 0.721 |
| Tryptophane | 0.722 | 0.504 | 0.507 | Cis-Aconitic acid | 0.272 | 0.279 | 0.329 |
| **Total** | **18.472** | **4.834** | **5.865** | Citric acid | 0.458 | 0.813 | 0.831 |
| **2. Fatty acid** |  |  |  | Isocitric acid | 0.196 | 0.478 | 0.537 |
| Capric acid | 0.473 | nd | nd | **Total** | **25.132** | **62.686** | **89.000** |
| Lauric acid | 0.478 | nd | nd | **4. Inorganic compounds** |  |  |  |
| Myristoleic acid | 0.150 | nd | nd | Na<sup>+</sup> | 58.87 | 83.65 | 85.05 |
| Myristic acid | 1.175 | 0.074 | 0.076 | NH<sub>4</sub><sup>+</sup> | 6.93 | 17.76 | 19.45 |
| Isopentadecylic acid | 0.092 | nd | nd | Mg<sup>2+</sup> | 29.24 | 27.52 | 27.95 |
| Isopalmitic acid | 0.114 | nd | nd | K<sup>+</sup> | 15.98 | 58.95 | 61.32 |
| Palmitoleic acid | 0.351 | 0.089 | 0.081 | Ca<sup>2+</sup> | 87.93 | 114.59 | 119.49 |
| Palmitic acid | 1.287 | 0.115 | 0.108 | Cl<sup>-</sup> | 43.34 | 120.21 | 116.32 |
| Linoleic acid | 0.404 | nd | nd | NO<sub>3</sub><sup>-</sup> | 489.67 | 398.43 | 384.51 |
| Oleic acid | 0.404 | nd | nd | SO<sub>4</sub><sup>2-</sup> | 5.41 | 130.15 | 128.64 |
| Stearic acid | 2.539 | 0.093 | 0.166 | PO<sub>4</sub><sup>2-</sup> | 10.66 | 4.74 | 4.18 |
| Arachidic acid | 1.076 | 0.071 | 0.074 | **Total** | **748.03** | **956.00** | **946.91** |

Method NSE was developed in this study; method TSE1 involves autoclaving at 121°C for 1 h; method TSE2 does not involve autoclaving. Soil components without supplements were derived from the sample, but different methods were used. Mean ± standard deviation; nd: not detected.

### Method validation based on isolation rate of uncultured or new taxonomic bacteria

To compare three methods: the newly developed method for isolation of previously uncultured soil bacteria using ISEM (intensive soil extract medium: simply new method), traditional soil extract culture medium (simply traditional method), and modified transwell culture method (simply modified method), 258, 243, and 252 pure colonies were isolated, respectively, from three soil samples as described in materials and methods. The ratio of uncultured bacterial isolates (previously uncultured strains) was significantly increased by 49% (126 isolates/total 258 isolates) for the new method, with values of 5% (11 isolates/total 243 isolates) for the traditional method and 15% (39 isolates/total 252 isolates) for the modified method (Fig. 1a; Tables S1-S3).

Taxonomic analysis also showed that this new method was much better than the two other methods in terms of new taxa isolation. In the new method, 142 new species candidates were found among isolates, including 13 genus level and 1 family level, showing 55% (142/258) efficiency compared to 13% (32/243) and 26% (65/252) for the traditional and modified methods, respectively (Tables S1-S3). For the isolation efficiency at the genus level or higher candidates, the new method showed a value of 5.4% (14/258), which is much higher than 0.0% (0/243) and 0.4% (1/252) obtained for the other two methods (Fig. 1b). The new method isolated a family level candidate, while the other two methods did not. Furthermore, the new method showed the highest ratio and largest number of new taxa candidates (at least species level) among uncultured isolates (75.4%: 95/126) compared to the other two methods, which showed values of 36.4% (4/11) and 48.7% (19/39), respectively (Fig. 1c).

In addition, our method is direct bacteria isolation from a soil suspension by using agar plates without an enrichment culture step, saving time and labour.

**FIG 1.**
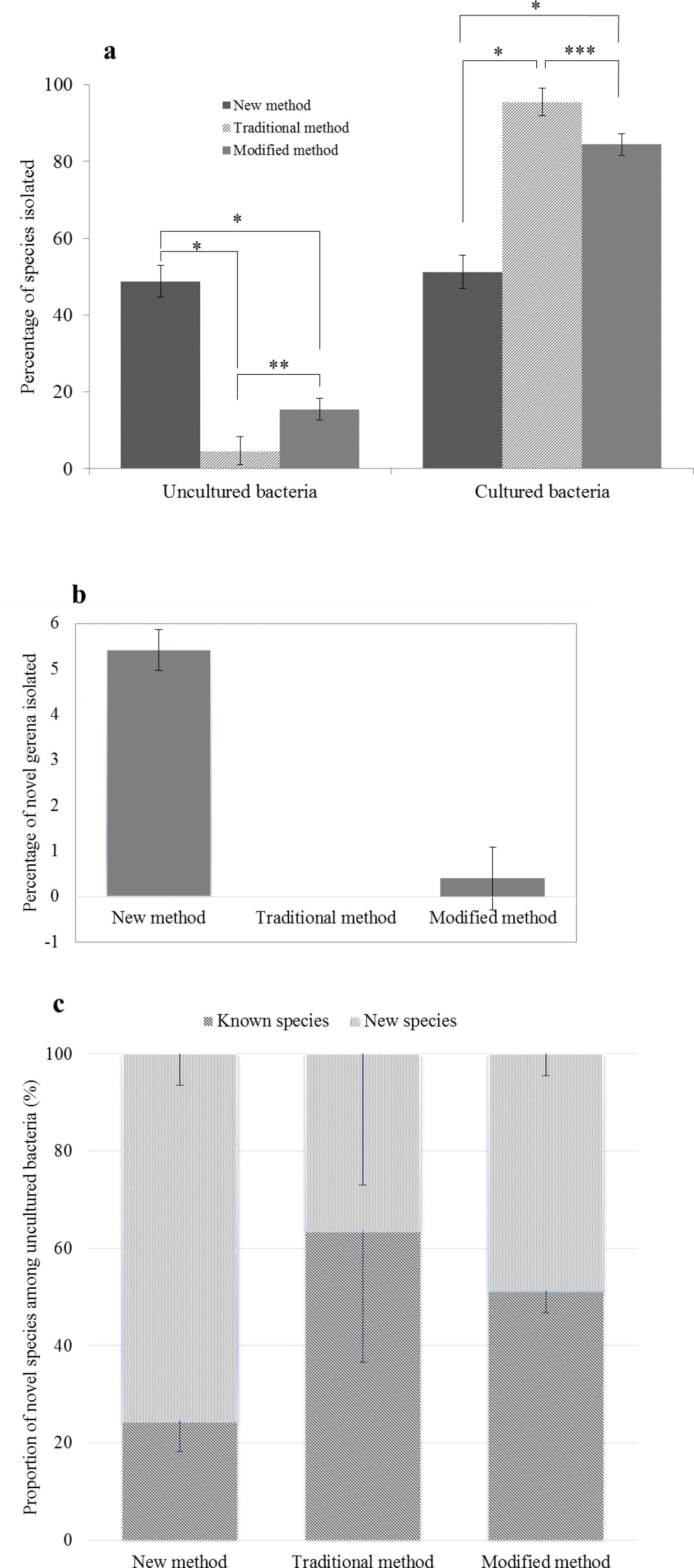
Proportion of uncultured bacteria and new taxonomic candidates isolated from soil samples by using ISEM (newly developed method), traditional soil extract culture medium and modified transwell culture method. (**a**) Percentage of uncultured species (126 individual species among the total 258; 11 of 243; 39 of 252, respectively). Points of significance compared to two paired samples for means t-test as indicated by * (P<0.005), ** (P<0.01), and *** (P<0.009). (**b**) Percentage of novel genera candidates (14/258; 0/243; 1/252, respectively). (**c**) Proportion of novel bacterial species and known species among 126, 11, and 39 previously uncultured species via the investigated methods, respectively.

### Method validation through taxonomic analysis

16S amplicon sequencing data as a next generation sequencing technology were analysed according to Chao1 at a 3% evolutionary distance and revealed the diversity of microbial community genomics in the soil samples, with 1744-2402 operational taxonomic units (Fig. S1a). Pyrosequencing analysis suggested that nine identified bacterial phyla commonly present in all soil samples were *Chloroflexi, Planctomycetes, Verrucomicrobia, Bacteroidetes, Gemmatimonadetes, Actinobacteria, Acidobacteria, Proteobacteria*, and *Parcubacteria*_OD1 (Fig. 2a). *Proteobacteria* and *Acidobacteria* were the most dominant phyla. Minor phyla (<1%) involved in 3.1% ETC were major lineages with the following isolated representatives: *Chlorobi, Elusimicrobia, Armatimonadetes, Firmicutes, Chlamydiae*, and *Tenericutes*; major lineages lacking representative: *Latescibacteria*, *Omnitrophica*, and *Hydrogenedentes*; candidate phyla: *Kazan, Gracilibacteria, Berkelbacteria*; other unidentified phyla (32). Additionally, the distribution and relative abundance of species identified by pyrosequencing in each soil sample were determined, showing that the species with more individuals would be less and vice versa (Fig. 2b). Phylogenetically, the new method achieved successful cultivation of strains from seven phyla among all bacteria present in the soils (Fig. 3): *Proteobacteria* (α, β, and γ) (46.9%), *Actinobacteria* (43.4%), *Bacteroidetes* (4.7%), *Firmicutes* (3.9%), *Acidobacteria* (0.4%), *Verrucomicrobia* (0.4%), and *Planctomycetes* (0.4%) (Fig. S1b). In contrast, the two other methods did not recover strains in three phyla: *Acidobacteria*, *Verrucomicrobia*, and *Planctomycetes*. Thus, the new method extended the taxonomic range of cultivation at the phylum level. Pyrosequencing analysis indicated that the three soil samples included 120 identified families, excluding 10% unclassified sequences, and 38, 22, and 29% of the identified families were recovered by the new, traditional, and modified methods, respectively (Fig. S2a). The isolates obtained using the new method represented 100 genera (86 known and 14 novel genera), which are compared with the 50 and 60 genera isolated using the traditional and modified methods, respectively (Fig. S2b). For the comparison at the species level, the new method independently cultivated soil bacteria as compared to the other two methods, as only one species among the 258 species overlapped with the traditional method (none with the modified method), while the two other methods showed 31 overlapping species with each other (Fig. S2c). Furthermore, uncultured isolates from the new method were distributed in 53 genera of six phyla, while the other two methods showed a very limited taxonomic distribution: 5 genera in two phyla and 20 genera in three phyla, respectively (Fig. S2d).

**FIG 2.**
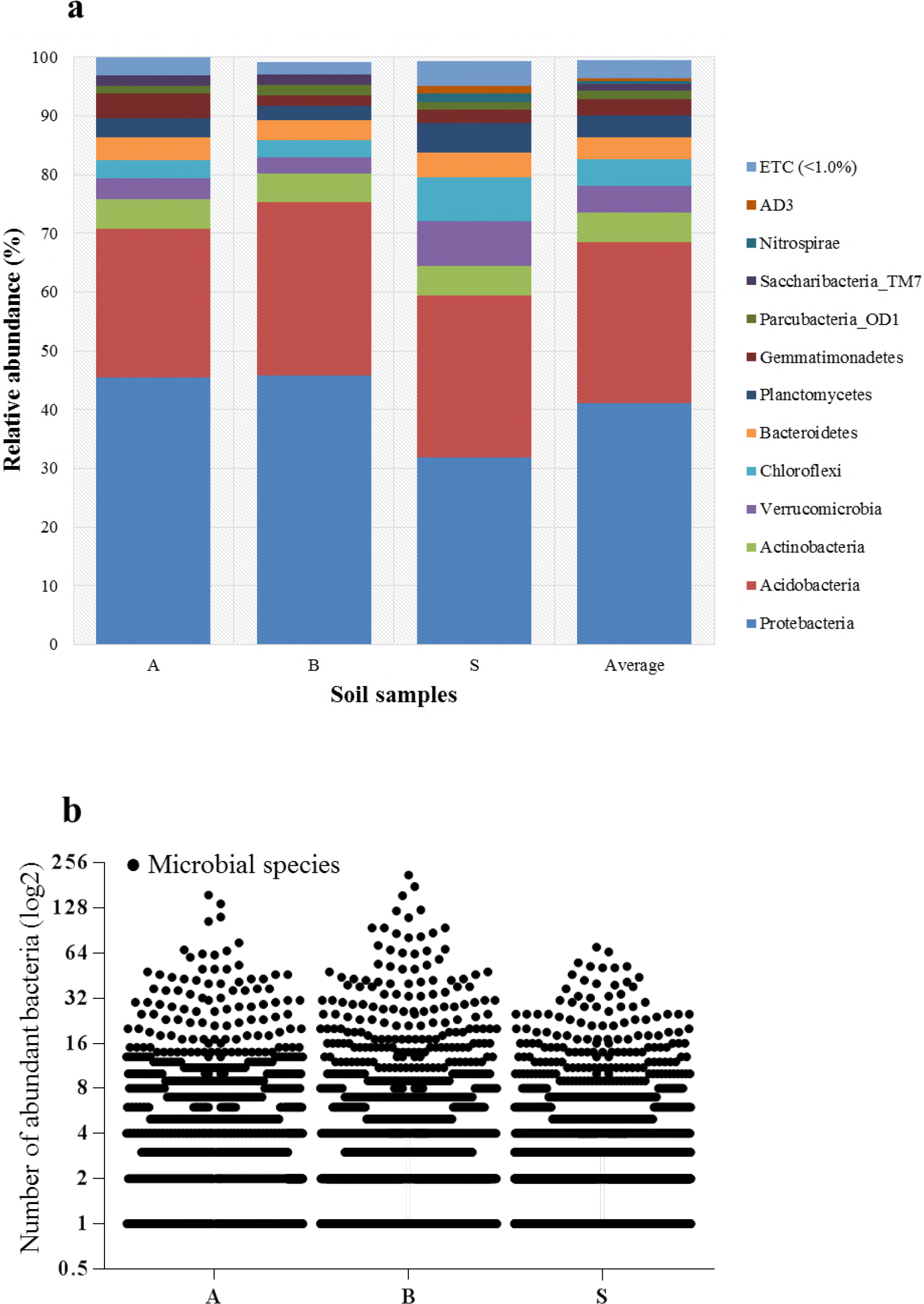
Abundance of bacteria in soil samples determined by pyrosequencing. **(a)** The abundance determined at a 1% ETC cut-off; minor phyla (<1%) present in each soil sample were *Chlorobi, Elusimicrobia, Latescibacteria, Armatimonadetes, Firmicutes, Kazan, Chlamydiae, Omnitrophica_OP3, Tenericutes, Gracilibacteria, Hydrogenedentes_NKB19, Berkelbacteria*, and other unknown phyla. **(b)** Distribution and relative abundance of identified species: approximately 1.2-1.3x10^6^ individual species detected in each soil sample.

**FIG 3.**
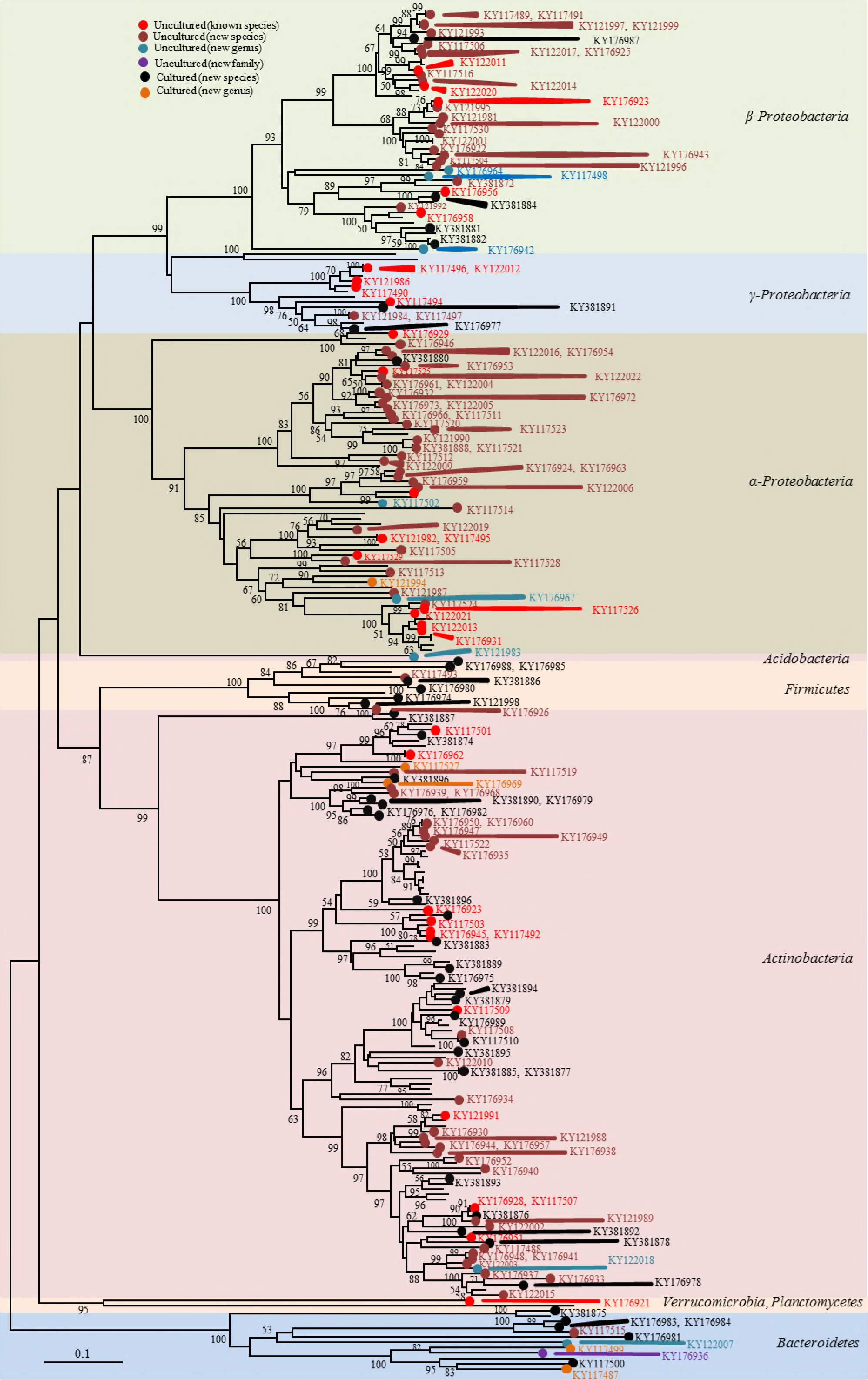
Network topology tree for microbial cultivation based on full-length 16S rRNA gene sequencing (n=258) showing seven phyla including: *Proteobacteria* (sub-phylum α, β, and γ), *Acidobacteria; Firmicutes, Actinobacteria, Verrucomicrobia, Planctomycetes, and Bacteroidetes*. Only bootstrap support values ≥50% are shown in the tree. Accession numbers for 16S rRNA gene sequences revealed close relationships with previously uncultured bacteria, and those cultured as new species and novel genera are shown in the tree.

### Method validation through cultivability of uncultured bacteria

We tested 7 different media for comparison: basic salts (BS as a negative control), BS plus micro-nutrients, BS plus vitamin B, BS plus D-amino acids, BS plus NSE, NSE only, and full ISEM (as a positive control). Here, 131 bacterial isolates (= 131 species) including 126 previously uncultured bacteria and 5 novel genus candidates (Table 2; S4), obtained using our developed culture method and three different soil samples, were used to determine the effectiveness of the various nutrient components by forming visible colonies on agar plates after streaking. While NSE and BS plus NSE showed 100% growth recovery (n = 131/131), BS plus D-amino acids showed a value of 8% (n = 11/131) and other components had a value of 0% (n = 0/131). Therefore, NSE and NSE-containing media can be useful as more effective media for isolating and subculturing uncultured soil bacteria than other media.

**Table 2.** **List of bacterial isolates used to evaluate new soil extract (NSE) and other media**

| Soil samples | Species level |  | Genus level |  | Family level | Total |
| --- | --- | --- | --- | --- | --- | --- |
|  | Previously uncultured novel species | Previously uncultured known species | Previously uncultured novel genus | Previously cultured novel genus | Previously uncultured novel family |  |
| A | 24 | 12 | 2 | 3 | - | **41** |
| B | 28 | 9 | 3 | 1 | - | **41** |
| S | 34 | 10 | 3 | 1 | 1 | **49** |
| Total | **86** | **31** | **8** | **5** | **1** | **131** |

## DISCUSSION

Soil contains necessary elements for living organisms. An aqueous soil extraction method was previously established (for example see ref. 36 or ATCC medium 654 or DSMZ medium 80) and remains widely used. Most mineral or organic ingredients such as ionic salts, vitamins, antibiotics, plant hormones, and plant-promoting growth factors, among others, can be dissolved in distilled water, while some organic components such as non-polar compounds cannot be sufficiently dissolved in distilled water or aqueous buffers. Thus, we used 80% methanol to overcome this problem and achieved higher concentrations and more different types of organic ingredients compared to using distilled water (Table 1). Although methanol and water are polar protic solvents that easily solubilize polar molecules, methanol is less polar than water based on their polarity values of 5.1 and 10.2, respectively. Therefore, methanol may more easily dissolve or extract a greater amount of hydrophobic or amphipathic molecules in soil than water. In contrast, based on their dielectric constants according to Harris (33), water (approximately 80) is more likely to dissolve inorganic compounds than methanol (approximately 30).

NSE contained 21 amino acids, which was greater than that obtained using water extraction methods (n=18) in this study (Table 1) and in previous studies: 16 amino acids were obtained using 6 N HCl (34) and 14 amino acids were obtained using MOPS buffer (31). Fatty acids contain a polar carboxylic group and non-polar hydrocarbon group of 4-36 carbons, making only short-chain fatty acids more or less water-soluble. Thus, a combination of methanol and water improves their dissolution. This led to significant differences in the total number and amount of fatty acids between methods NSE and TSE1 (P<0.003), but the total fatty acids obtained for the two comparative methods (method TSE1, and method TSE2) were similar each other (*P*>0.9) (Table 1). Organic acids are widely present in soil (31) and low-molecular weight organic acids are typically miscible in water. Thus, the three extraction methods were relatively effective. Although methanol extracted lower concentrations than water, more diverse organic acids were extracted, increasing the spectrum of either carbon or electron donors/acceptors for microorganisms. Because inorganic substances typically dissolve well in water and even in pure methanol (35), 80% methanol can extract large amounts of inorganic compounds (5.41-489.67 mg/L) from soil, although lower amounts than the two water extraction methods. We supposed that the components present only in NSE or much higher concentrated in NSE than in other two TSEs might help to stimulate growth of uncultured bacteria. Those were alanine, valine, leucine, isoleucine, pipecolic acid, serine, threonine, and tyrosine in amino acid; capric acid, lauric acid, myristoleic acid, myristic acid, isopentadecylic acid, isopalmitic acid, palmitic acid, linoleic acid, oleic acid, stearic acid, arachidic acid behenic acid, nervonic acid, and lignoceric acid in fatty acid; lactic acid, glycolic acid, 2-hydroxybutyric acid, fumaric acid, and a-ketoglutaric acid in organic acid. Overall, NSE was superior compared to the two traditional soil extracts (TSEs) and other extraction methods because greater concentrations and types of LMWOS were obtained, which are required to support most soil bacteria including uncultured bacteria.

Although an enhanced medium (traditional soil extract culture medium) derived from soil extract (TSE) containing yeast extract, tryptone, and salts was designed to support various soil bacteria (36), necessary elements for many uncultured bacteria may be absent, so that most isolates (~96%) seemed to belong to previously cultured groups (Fig. 1a; Table S2). Additionally, the modified method with R2A, a complex efficient artificial media for cultivating heterotrophic bacteria including fast-growing and slow-growing bacteria (37), showed better results and isolated a greater number of uncultured bacteria and new taxa candidates than the traditional method did (Fig. 1a; Fig. 2a, b; Table S3). Although this method is better than the traditional method, the isolation step shows limited recovery of various enriched uncultured soil bacteria. Particularly, ISEM developed in this study allowed cultivation of large numbers of isolates of various uncultured bacteria and new taxa candidates compared to the numbers obtained using the other two methods. Thus, the new method more effectively isolated uncultured bacteria and new bacterial taxa present in soil. The most important aspect of this new method (ISEM) is to include NSE that is described precedingly. As mentioned in the result, the direct isolation from a soil suspension using ISEM agar plates can be an effective way because, in fact, during the enrichment step, fast-growing microorganisms may overcome slow growing bacteria, leading to reduced diversity of bacterial species.

Among recently introduced methods, Kakumanu & Williams (2012) (38) developed a soil diffusion system and found uncultured bacteria in the phyla *Proteobacteria, Bacteroidetes, Verrucomicrobia, Planctomycetes*, and OP10, but only 8 uncultured bacteria at the species level. Furthermore, the study suggested conducting only enrichment culture, with no method for isolation of pure bacterial strains. Another study found that 27% of species belonged to 20 unnamed family-level groupings among 350 isolates (39), but these species were isolated from many different artificial media, not including soil extracts, and the effectiveness of each media on the cultivability of uncultured soil bacteria was not determined. Various culture methods for cultivating uncultured soil bacteria have been developed: modification of growth media, modification of growth conditions, community culture, coculture, trans-well plates with membranes, micromanipulator, optical tweezers, laser microdissection, high throughput microbioreactor, simulated natural environments using diffusion chambers, single cell encapsulation combined with flow cytometry, multiwell microbial culture chip or iChip, and entrapped gelating agent coated with polymer (30). However, these methods exhibit low isolation efficiency of uncultured bacteria or lack strategies for continuous pure culture. The new method developed in this study showed high isolation efficiency (49%), a 100% recovery rate of uncultured soil bacteria, and easier application in laboratories compared to most previously developed methods.

Although *Acidobacteria* were secondly abundant phylum in pyrosequencing analysis for three soil samples, only one isolate was obtained through our new method since the medium (ISEM) might be not acidic (pH 6.8). As a future work, if using ISEM with low pH or low temperature, more uncultured or novel acidobacteria or other bacteria may successfully be isolated.

In summary, our new method showed a much higher isolation rate of new taxa candidates among uncultured isolates and greater isolation rate of uncultured soil bacteria and new taxa candidates than traditional and modified methods tested in this study. Additionally, isolation was simpler and did not require enrichment culture, and could be directly subcultured to obtain more uncultured bacterial pure masses for further experiments. Further, the variety of uncultured soil bacteria or new taxa candidates can be extended with ISEM by altering the cultivation conditions such as incubation temperature, pH, salt concentration, anaerobic conditions or using various soil samples and other samples.

## MATERIALS AND METHODS

### Soil sample used for making soil extracts

Rhizosphere soil where *Robiniapseudoacacia* L. dominated was collected at Kyonggi University (154-42 Gwanggyosan-ro, Iui-dong, Yeongtong-gu, Suwon, Gyeonggi-do, South Korea; 37°30′04″ N, 127°03′58″ E) during June 2016. Fresh soil was dried at room temperature for 48 h by spreading soil sample on surface of aluminum foil and using an air conditioner (25~30°C), and then any plant debris, gravels and rocks were removed by a 0.2 mm sieve. To determine physicochemical properties of soil, the soil was dried at 110°C for 24 h, and cooled at room temperature. Soil contained approximately 78% sand, 17% silt and 5% clay. Its pH was ~5.7 that was defined directly from fresh soil.

### Preparing new soil extract (NSE) in 1 L medium

Approximately, 1000g of the dry soil sieve at room temperature was divided into two equal parts (500g each) in a 2-L flask and then mixed with 1.3 L of 80% methanol (#494291, grade methanol, Sigma Aldrich, St. Louis, MO, USA) in deionized water and shaken at 150 rpm overnight at room temperature (below 25°C). After allowing settling for 30 min, the supernatant was transferred to a new flask. Next, 1.3 L of 80% methanol was added to the soil and mixed well for 1 h. The two supernatants were combined and filtered through Whatman^TM^ paper (#1001-150, ϕ 150 mm, GE Healthcare, Little Chalfont, UK). Methanol was removed by a general rotary evaporator (~40°C). The NSE was adjusted to a final volume of 200 mL with deionized water and sterilized through a 0.22μm filter membrane composed of nitrocellulose (#GSWP04700, Merck Millipore Ltd., Billerica, MA, USA) using a vacuum pump. The pure supernatant was passed through via a 0.2μm polyvinylidene fluoride membrane (#WHA67791302, Sigma Aldrich) if necessary to ensure that no bacteria were in the soil extract, and pure NSE was stored in a dark Schott Duran bottle at 4°C and used within one week.

### Medium for isolation of soil bacteria

The medium containing 0.23 g KH_2_PO_4_, 0.23 g K_2_HPO_4_, 0.23g MgSO_4_·7H_2_O, 0.33 g NH_4_NO_3_, 0.25 g NaHCO_3_ as a group of mineral salts, and 15 g agar in 1 L water was sterilized at 12°C for 15 min. To the medium was added 5 mg various D-amino acids (D-valine, D-methionine, D-leucine, D-phenylalanine, D-threonine, and D-tryptophan), 1 mL vitamin B (vitamin stock solution containing 50 mg each thiamine hydrochloride, riboflavin, niacin, pyridoxine HCl, inositol, calcium pantothenate, and β-aminobenzoic acid and 25 mg biotin in 100 mL distilled water, sterilized through a 0.2μm syringe filter, stored at 4°C in dark Schott Duran and used within one month), 0.2 L of NSE, 2 mL of selenite-tungstate solution (40) (composition in 1 L distilled water: 0. 5g NaOH, 3 mg Na_2_SeO_3_.5H_2_O, 4 mg Na_2_WO_4_.2H_2_O; the solution was filter-sterilized, stored at 4°C and used within 1 month), and 2 mL of trace element SL-10 (41) (ingredient contained 10 mL of HCl (25%, v/v); 1.5 g of FeCl_2_4H_2_O; 70 mg of ZnCl_2_; 100 mg of MnCl_2_4H_2_O; 6 mg of H_3_BO_3_; 190 mg of CoCl_2_·6H_2_O; 2 mg of CuCl_2_·2H_2_O; 24 mg of NiCl_2_·6H_2_O; 36 mg of Na_2_MoO_4_·2H_2_O in a final volume of 1L; then this solution was passed through a 0.2μm filter, added directly in the medium after autoclaving). The final volume was 1L and pH was 6.8±0.2. The complex medium was named as intensive soil extract medium (ISEM). The medium should be prepared freshly and used within one week. In this study, we used 150×20 mm petri dishes (SPL Life Science Co., Ltd., Gyeonggi-do, Korea). The larger dish allows for increased separation of colonies at high dilution concentrations during isolation.

### Preparation of various media to recover previously uncultured soil bacterial isolates

We used multiple combinations to find a best growth medium as described above to identify the most important elements for supporting the growth of previously uncultured soil bacteria. Several examinations were carried out based on the strains obtained in this study as follows (i): basic salts (BS) only as a negative control, (ii): BS with selenite-tungstate solution and SL-10; (iii): BS plus D-amino acids, (iv): BS with vitamin B, (v): BS with NSE, (vi): NSE only; (vii): mixture of all components (ISEM) as a positive control, 15 g agar (#A7049, Sigma-Aldrich) was treated several times with distilled water to discard any trace nutrients or elements, and then added to each medium. Agar plates were incubated at 25°C for 4 weeks under aerobic conditions.

### Soil sampling sites and preparation of soil samples

Three soil samples were acquired in South Korea in June 2016, including Ansan (sample A) (Il-dong, Sangnok-gu, Ansan, Gyeonggi-do: 37°17’58”N & 126°53’57 E), Suwon (sample B) (Buksu-dong, Paldal-gu, Suwon, Gyeonggi-do: 37°16’42”N & 127°00’17” E), and Seoul (sample S) (Itaewon-ro, Yongsan-gu, Seoul: 37°31’06” N, 127°01’04” E). For each sample, approximately10 g soil from ten different locations within a 150-m diameter were collected and mixed well. The sample was passed through 0.1-mm sieve and isolated/enriched directly using three methods. A 25-g sieved soil sample was mixed with 250 mL sterile saline (0.9% NaCl, w/v), stirred for 15 min, and allowed to separate between suspension and sediment before use.

### Newly developed method for isolation of previously uncultured soil bacteria

First, 100 μL of each the dilution of soil suspension was spread onto three agar plates of ISEM (to ensure uniformly distributed suspension on the surface of the medium, 100 μL of each the dilution plus 100 μL of ISEM liquid is recommended). These agar plates were incubated at 25°C for 6 weeks. A few colonies appeared after one week of incubation. The number of directly observable colonies was increased after 2 weeks, and tiny colonies were picked up and streaked onto fresh ISEM until morphologically pure colonies were obtained. Cells on fresh ISEM typically require at least one week growing. Uncultured bacteria generally showed weak growth, and thus in some cases pure colonies were activated in ISEM broth in a shaking incubator at 25°C, 150 rpm for 1-2 weeks before transferring onto the agar plate.

### Traditional soil extract culture medium

Approximately 1000 g air-dried soil in 1.3 L of deionized water was autoclaved at 121°C for 1 h and allowed to cool. The supernatant was filtered through Whatman^TM^ paper before centrifugation in a 500-mL bottle at 5009 ×g for 30 min at room temperature. One litre of the supernatant (TSE) was obtained. Soil extract agar was enhanced by supplementation (42) with 0.04% K_2_HPO_4_, 0.005% MgSO_4_.7H_2_O, 0.01% NaCl, 0.001% FeCl_3_, 0.05% tryptone, 0.05% yeast extract, and 1.5% agar in 1 L of soil extract liquid with a final pH of 6.8. Next, 100 μL of each dilution of three soil samples was dispersed onto three soil extract agar plates and cultivated at 25°C for 6 weeks. Colonies were re-streaked until pure colonies were obtained.

### Modified transwell culture method

Transwell plate system (#35006, SPLInsert™ Hanging, SPL Life Sciences) was used to enrich bacterial community, especially for uncultured soil bacteria, from soil samples, which contains 6-inserts with 6-wells in a plate. An insert has two different sized frames (uppper: 28 mm outer, and 26.65 mm inner; lower: 26.6 mm outer, and 23.3 mm inner); 28 mm height, and 4.52 cm^2^ of area of growth for each insert. The lower frame is covered with a 0.4μm polycarbonate membrane. Its membrane specification is 25 mm diameter, and 7~10 p,m thickness. Approximately 3 g of soil sample was added to a transwell plate, and then 3 mL R2A medium (#MB-R2230, MB Cell, Los Angeles, CA, USA; 3.15 g of the powder in 1 L distilled water) was supplemented into the soil-containing wells and then put the insert on the wet soil. Next, 100 μL of the suspension and 1 mL R2A medium was inoculated into the insert. The transwell culture system was covered with parafilm to prevent evaporation. The system was shaken at 120 r.p.m and 25°C for 4 weeks. Seven-fold dilutions of the culture enriched were established in R2A broth medium; 100 μL of each dilution was spread onto three R2A agar plates and incubated at 25°C for 6 weeks. Colonies were subcultured on R2A medium to obtain individual colonies (Table S5).

### Identification of 16S rRNA gene sequences and accession numbers of 16S rRNA gene sequences

Near-full-length 16S rRNA sequences were identified and similarity to valid species was calculated using the EzTaxon Database Update (http://www.ezbiocloud.net/eztaxon) for comparison with published uncultured gene sequences via the nucleotide BLAST in NCBI: http://www.ncbi.nlm.nih.gov/. In this study, based on full 16S rRNA similarity to valid published species, with four temporary divisions, candidates of novel species were defined by comparison of 16S rRNA similarity at the threshold of 98.7% (2), 95.3-90.0% novel genus level (3), and novel family level at off limit lower than 90.0%. Candidates of uncultured or novel species or new genera or family sequences were submitted to GenBank database and are listed in the Supplementary Tables; isolates without accession numbers are available upon request.

### DNA extraction from soil

Using FastDNA^®^ SPIN Kit for Soil (#116560-200, MP Biomedicals), soil DNA from 0.5 g of fresh soil was extracted and purified by following the instruction’s guide. DNA quality was checked through 1.2% agarose gel electrophoresis in 0.5× TAE buffer and DNA concentration was determined via MaestroNano spectrophotometer (Mastrogen). Then DNA samples were held at −20°C until use.

### PCR amplification and pyrosequencing

Pure isolated DNA soil samples were applied subjected to amplification of the target V1 to V3 regions located in the 16S rRNA gene by PCR using the barcoding primers 27F 5′-CCTATCCCCTGTGTGCCTTGGCAGTC-TCAG-AC-GAGTTTGATCMTGGCTCAG-3′ and 518R 5′-CCATCTCATCCCTGCGTGTCTCCGAC-TCAG-X-ACWTTACCGCGGCTGCTGG-3′; ‘X’ directs the unique barcode for each subject) (http://oklbb.ezbiocloud.net/content/1001). The reaction was conducted as follows: initial denaturation at 95°C for 5 min, followed by 30 cycles of (denaturation at 95°C for 30 sec, annealing at 55°C for 30 sec, and extension at 72°C for 30 sec), and final elongation at 72°C for 5 min. Next, the amplicons were evaluated by 2% agarose gel electrophoresis and observed with a Gel Doc system (Bio-Rad, Hercules, CA, USA). A QIA quick PCR purification kit (#28106, Qiagen, Hilden, Germany) was used to purify the PCR products. An Ampure beads kit (Agencourt Bioscience, Beverly, MA, USA) was used to enhance the quality of the sample and remove non-target products following the manufacturer’s instructions. Quality and target size were estimated with a Bioanalyzer 2100 (Agilent, Santa Clara, CA, USA) using a DNA 7500 chip. Next, PCR products were mixed by emulsion PCR and deposited on Picotiter plates. Target sequencing was conducted with a GS Junior Sequencing system (Roche, Basel, Switzerland) according to the manufacturer’s instructions.

### Analysis of pyrosequencing data

Pyrosequencing results were analysed as follows. Unique barcodes for each amplicon as a standard were sorted from distinctive samples and readings were obtained. Removal of either these non-target sequences including the barcode, linker, and primers or more than two ambiguous nucleotides, low-quality score of less than 25 through trimmomatic 0.321 (43), and reads shorter than 300 bp from the original sequencing reads. Additionally, the Bellerophone method was used to discard chimeric sequences, and then a full sequence was compared both in the forward and reverse directions via BLASTN (44). The similarity of each full sequence to valid published type strains was determined using the EzTaxon-e database (http://eztaxon-e.ezbiocloud.net) or uncultured bacterium clone in GenBank database (https://blast.ncbi.nlm.nih.gov). Chao1 estimation at a 3% distance (45) and Shannon diversity index (46) were used to confirm the level of richness and diversity of each sample. Phylogenetic analysis of microbial communities was estimated via the Fast UniFrac (47) combined with principle coordinate analysis. Finally, XOR analysis of CLcommunity program (Chunlab Inc., Seoul, Korea) was used to compare the number of operational taxonomic units among samples.

### Determining and comparison of soil extract ingredients

To determine the impacts of different ingredients, the rhizosphere soil (*Robinia pseudoacacia* L.) at Kyonggi University was collected and prepared in three methods: the first as NSE; the second as TSE1 was autoclaved at 121°C for an hour, allowed cool and settling, next the supernatant was passed through a 0.22μm filter nitrocellulose membrane using a vacuum pump, followed by a rotary evaporator bath at 40°C to reduce water to 200 mL; the third method as TSE2 was similar to TSE1 except sterilization (at 121°C) was replaced with shaking overnight at room temperature to compare variations in soil components at high temperature. Three samples obtained from the supernatant after soil extraction methods were lyophilized at −54°C to produce 6.36 g powder (NSE), 4.65 g for TSE1, and 4.73 g for TSE2. These powders were stored at 4°C until analysis. For inorganic composition and carbohydrates, soils were prepared and analysed directly from a final volume of 1 L of deionized water without adding any substrates. Experiments were three repeated to ensure accuracy, and chemical data were prepared and analysed individually in triplicate.

### Sample preparation for simultaneous profiling analysis of amino acids, organic acids, and fatty acids in soil extract

Amino acids (AAs), organic acids (OAs), and fatty acids (FAs) were simultaneously profiled in soil samples as their ethoxycarbonylation (EOC), methoximation (MO), and *tert*-butyldimethylsilyl (TBDMS) derivatives as described previously (48, 49). Briefly, 2.5 mg soil extract was dissolved in distilled water containing 0.1 μg of norvaline, 3,4-dimethoxybenzoic acid, and pentadecanoic acid as internal standards. The solution pH was adjusted to ≥12 with 5.0 M sodium hydroxide and mixed with dichloromethane (2.0 mL) containing 40 μL ethyl chloroformate, which was converted to the EOC derivative. This was converted to the MO derivative via a reaction with methoxyamine hydrochloride at 60°C for 60 min. The aqueous phase as sequential EOC/MO derivatives was acidified (pH ≤ 2.0 with 10% sulphuric acid), saturated with sodium chloride, and extracted with diethyl ether (3 mL×2). The extracts were evaporated to dryness using a gentle nitrogen stream. Dry residues containing AAs, OAs, and FAs were reacted at 60°C for 30 min with TEA (5 μL), toluene (15 μL), and *N*-methyl-*N*-(tert-butyldimethylsilyl)trifluoroacetamide (20 μL) to form the TBDMS derivative. All samples were prepared individually in triplicate and examined directly by gas chromatography-mass spectrometry (GC-MS) in selected ion monitoring (SIM) mode.

### Inorganic ingredients

The cations Li^+^, Na^+^, Mg^2+^, K^+^, Ca^2+^, and NH_4_^+^ were detected using a Dionex ICS3000 (Sunnyvale, CA, USA) with an Ionpac CS12A column (4×250 mm/Dionex) and detector (suppressed conductivity, CSRS URTRA (4 mm), recycle mode). Oven temperature was 30°C, injection volume was 25 μL, samples were eluted with 20 mM methanesulfonic acid at flow rate 1 mL/min, and run time was 20 min according to the IonPac^®^ CS12Amanual (Thermo Fisher Scientific). A Dionex ICS3000 was also used to detect the anions F^−^, Cl^−^, Br^−^, NO_2_^−^, NO_3_^−^, SO_4_^2−^, and PO_4_^2−^ with standards, and the column was an Ionpac AS20 (4×250 mm, Dionex);the detector was a suppressed conductivity ASRS URTRA II (4mm), recycle mode. Gradient elution was conducted for 0-8 min (12 mM KOH), 8-12 min (30 mM KOH), 12-17 min (30 mM KOH), 17-18 min (12 mM KOH), and 18-20 min (12 mM KOH) at a flow rate 1 mL/min.The oven temperature was 30°C and injection volume was 25 μL. All processes were conducted as described in the IonPac®AS20 Anion Exchange Column product manual (Thermo Fisher Scientific).

## ACKNOWLEDGEMENTS

This research was supported by the Basic Science Research Program through the National Research Foundation of Korea (NRF) funded by the Ministry of Education (2016R1D1A1A09916982).

---

**FIG S1.** Relation between sequencing reads and OTUs of three soil samples analysed by Chao1 at the 3% evolutionary distance (a); number of genera in each phylum isolated from three methods (b).

**FIG S2.** Comparison of three culture methods by isolated families (a); genera (b); overlapping species among isolates obtained from three different methods (c), and number of genera in each phylum, previously uncultured, isolated from three methods (d).

**Table S1. List of isolates obtained by improved medium (new method).** Red color: known species; Green: new species candidate; Blue: new genus candidate; Purple: new family candidate.

**Table S2. List of isolates obtained by traditional method** Red color: Known species; Green: new species candidate.

**Table S3. List of isolates obtained by modified method** Red color: known species; Green: new species candidate; Blue: new genus candidate.

**Table S4. List of bacterial isolates used to evaluate new soil extract (NSE) and two traditional extracts (TSEs)**

**Table S5. Total colonies counted and picked up on dilution plates of soil sample suspensions (A, B and S) according to three culture methods**

